# Enhanced Hi-C Capture Analysis reveals complex regulatory architecture at the *PICALM_EED* locus for Alzheimer’s disease

**DOI:** 10.64898/2026.02.14.705927

**Authors:** Luciana Bertholim-Nasciben, Liyong Wang, Wanying Xu, Aura Ramirez, Sofia Moura, Leina Lu, Xiaoxiao Liu, Farid Rajabli, Katrina Celis, Marla. Gearing, David A. Bennett, Sandra. Weintraub, Changiz Geula, Theresa Schuck, Karen Nuytemans, William K. Scott, Derek M. Dykxhoorn, Margaret A. Pericak-Vance, Juan I. Young, Anthony J. Griswold, Fulai Jin, Jeffery M. Vance

## Abstract

**Objective:** Both the phosphatidylinositol binding clathrin assembly protein gene (*PICALM)* and the embryonic ectoderm development gene (*EED)* have been implicated as causal genes driving a genome-wide association for Alzheimer disease (AD) risk. We employed a new virtual approach using genome-wide chromatin interactions (Hi-C) called enhanced Hi-C Capture Analysis (eHiCA) to identify the genes and regulatory regions that are driving this important AD risk association.

**Methods:** Hi-C data from the frontal cortex of eight AD patients, as well as inducible pluripotent stem cell-derived microglia and spheroids of AD and control patients were used. We applied 14 eHiCA baits each containing a GWAS SNP to identify the *cis* regulatory interactions in this GWAS locus at a 5kb resolution.

**Results:** The baits derived from the GWAS associated haplotype primarily interacted with the PICALM promoter and the large cis-regulatory elements cluster (CREe) lying upstream of the *EED* promoter. The *EED* promoter interacts with *PICALM* gene body and promoter region but not directly with the associated risk haplotype. Although the AD-associated variants segregate together as a haplotype in the population, each bait exhibited distinct functional chromatin interactions.

**Interpretation:** The *PICALM* gene is the primary driver of the association in microglia along with the CREe locus. Different SNPs in a segregating haplotype can display different physical Hi-C interactions. This study demonstrates that eHiCA can help resolve the casual genes driving complex GWAS associations, opening new pathways to study Alzheimer disease and other disorders.

## Introduction

Identifying the casual target gene(s) driving a GWAS association has been a challenge limiting the interpretability of these studies. Several factors have contributed to this issue. First, non-coding regions often function as cis-regulatory elements (CREs) that, in the three-dimensional (3D) genome space, can interact with and regulate distant genomic sites. Second, associated SNPs usually lie in a haplotype of linked genetic variants segregating together with only a single or small number conferring risk. Finally, the functional impact of non-coding variants are both cell-type and cell-state specific.

The spatial DNA arrangement is an important epigenomic feature influencing interactions between promoters and CREs (Schoenfelder & Fraser, 2019). To facilitate mapping of the cis-regulatory architecture, we have applied a genome-wide chromatin interaction analysis, Hi-C (High-throughput chromosome conformation capture). To enhance the resolution of this Hi-C analysis, we utilized *DeepLoop*, which reduces technical bias and improves signal-to-noise ratio (Zhang et al., 2022). To facilitate focused studies centered on specific regions of interest, we developed a virtual variation using Hi-C, termed <u>e</u>nhanced <u>Hi</u>-C <u>C</u>apture <u>A</u>nalysis (eHiCA), which leverages enhanced Hi-C data from *DeepLoop* using a5kb bait to capture its’ focused interactions with other genomic regions. The 5 kb baits are smaller than the average haplotype (The International HapMap Consortium, 2005) enabling interrogation of chromatin loop interactions across multiple SNPs within a haplotype. When coupled with other -omics data, such as chromatin accessibility (ATAC-seq) and transcriptome data (RNA-seq), this approach provides a framework to delineate the potential causal variant(s) among a complex haplotypes and link these to effector target gene(s) that drive the GWAS association.

Specifically, we investigated the AD locus (Ando et al., 2022; Bellenguez et al., 2022) on chromosome 11q14.2 lying between two biologically plausible candidates, the phosphatidylinositol binding clathrin-assembly protein (*PICALM)* and the Embryonic Ectoderm Development *(EED)* genes (Ando et al., 2022; Kozlova et al., 2025; Moreau et al., 2014; Xiao et al., 2012; Zhao et al., 2015). Over 200 SNPs are associated with late-onset AD (LOAD) at this locus (p < 5 ×10^−8^), spanning a >240 kb region (Bellenguez et al., 2022). The most significant SNP (rs3851179) lies approximately 88 kb and 87 kb away from the canonical transcription start sites (TSS) of *PICALM* and *EED*, respectively. *PICALM* facilitates clathrin-mediated endocytosis - crucial for internalizing amyloid precursor protein (APP), regulating β-secretase cleavage, and modulating Aβ uptake or clearance in neurons and glia (Xiao et al., 2012; Zhao et al., 2015). *EED*, is an core component of the Polycomb Repressive Complex 2 (PRC2), binds trimethylated H3K27 marks to maintain transcriptional repression, thereby regulating gene expression programs fundamental for neural progenitor cell proliferation, differentiation, and cortical neurogenesis (Ueda et al., 2016). Given their mechanistic relevance, the Alzheimer’s disease Sequencing Project (ADSP) Gene Verification Committee lists *EED* as the potential susceptibility gene of the chr11q14.2 locus, and *PICALM* coding variants as causal elements. However, the exact genetic basis underlying the GWAS association remains unclear.

Herein, we integrated eHiCA using bulk Hi-C data with our Single-nucleus ATAC-seq (snATAC-seq) data (Celis et al., 2023; Griswold et al., 2021) in brain samples from individuals with AD to comprehensively map the regulatory architecture of the *PICALM/EED* locus and explore the causal AD association. To further delineate the cell-type specific variant-mediated regulatory architecture across this locus, we applied this integrative framework to frontal cortex, iPSC-derived microglia and neural spheroids derived from AD and non-cognitively impaired (NCI) patients.

## Methods

### Brain samples and iPSC-derived microglia and neurospheroids

Frozen frontal cortex from Brodmann area 9 from eight individuals were analyzed (Supplementary Table 1) (Celis et al., 2023; Griswold et al., 2021). All autopsy samples had a confirmed neuropathological diagnosis of AD. iPSC-derived microglia cells and neural spheroids from AD and NCI patients were generated and cultured in the Hussman Institute for Human Genomics iPSC Core facility using previously published protocols (Moura et al., 2025; Ramirez et al., 2025). All samples were acquired with informed consent for research use and approved by the institutional review board of each participating institution.

### Hi-C library preparation and sequencing

*In situ* Hi-C libraries were prepared using a protocol adapted from Rao et al (2014). Briefly, frozen tissue (≈100 mg) or iPSC-derived cells (1∼2 million) were crosslinked with formaldehyde (1% final concentration). The crosslink reaction was quenched with glycine (0.2M final concentration). After cell lysis, nuclei were permeabilized with 0.5% SDS and quenched with 1% Triton X-100. DNA was digested *in situ* overnight using *Mbo I* (New England Biolab). After restriction digestion, the DNA ends were filled in and labeled *in situ* with biotinylated-dATP (Active Motif) before proximity ligation. Then, crosslinks were reversed and the ligated DNA was purified. The purified DNA was sheared using the Covaris LE220 (Covaris, Woburn, MA). DNA fragments in the range of 300-500 bp were enriched with AmPure XP beads (Beckman Coulter). Ligation junctions with biotin labels were pulled down using streptavidin beads (Invitrogen) and prepared for Illumina sequencing using 8 cycles of PCR amplification. For each library, 340∼860 million paired-end reads of 150 bp length were generated using the Illumina Novaseq X 25B.

### Chromatin loop calling with enhanced Hi-C data

We first performed primary Hi-C data analysis with *HiCorr (*Lu et al., 2020) and *DeepLoop (*Zhang et al., 2022) pipelines. To that end, sequencing data were processed using BWA ^19^ to map each read end separately to the hg38 reference genome. Duplicate and non-uniquely mapped reads were removed. Two SAM files for each end were merged into a single file. Read pairs were assigned to restriction enzyme *Mbo I* fragments and read pairs located within same fragment being discarded. The *Mbo I* fragments are further assigned to 5kb bins and the Hi-C data was eventually represented as two-dimensional contact matrices of 5kb anchors. Next, we used the *HiCorr* pipeline for Hi-C bias correction. *HiCorr* explicitly corrects distance effects in a joint function with other biases followed by an implicit bias-correction step to balance the “visibility” of all 5kb bins in the genome. Notably, *HiCorr* outputs matrices of observed/expected ratios for chromatin interaction profiling. Finally, we applied *LoopEnhance* function in the *DeepLoop* package to the *HiCorr*-corrected contact matrices. These enhanced contact matrices were used for eHiCA analysis.

### Enhanced Hi-C Capture Analysis (eHiCA)

Like virtual 4C plots, eHiCA visualizes interacting signals from any bait-of-interest with the signals extracted from *DeepLoop*-enhanced Hi-C contact matrices. As a result, the plotted signals are enhanced observed *versus* expected ratios. To perform eHiCA, a 5kb region-of-interest (the “bait”) is derived from a genome-wide significant-associated GWAS haplotype with the investigated SNP lying in the center of the bait. Next, the enhanced Hi-C contact matrix data involving the “bait” are extracted, capturing the chromatin interactions between the “bait” and the genes/regions with which it physically interacts. We show the interacting signal +/-2 Mb surrounding this bait depicting the loop strength of the interactions using gradient heatmaps.

### Chromatin accessibility and RNA expression data

Chromatin accessibility and transcriptomic profiles were obtained from both single-nucleus and bulk sequencing datasets. SnATAC-seq and single-nucleus RNA-seq (snRNA-seq) from frozen frontal cortex (Brodmann area 9) were generated as previously described (Celis et al., 2023; Griswold et al., 2021). To complement brain-derived single-nucleus data, we also utilized bulk RNA-seq and bulk ATAC-seq generated from induced pluripotent stem cell–derived microglia (iMGLs) as described in (Moura et al., 2025).

### Co-localization of GWAS SNPs with ATAC-peaks

GWAS significant SNPs for the *PICALM/EED* locus (Bellenguez et al., 2022) were colocalized with in-house snATAC-seq (single nuclei Assay for Transposase-Accessible Chromatin using sequencing) data from the frontal cortex (Celis et al., 2023).

## Results

### Regulatory potential of SNPs associated with AD at the *PICALM_EED* locus

Multiple LOAD GWAS have identified genome-wide significant signals in chromosome 11 spanning a >240 kb region overlapping *PICALM* and the intergenic region between *PICALM* and *EED (Bellenguez et al*., *2022)*. In addition to the GWAS sentinel SNP rs3851179, among the 216 SNPs at this locus associated with LOAD at genome-wide significance threshold (*P* < 5 ×10^−8^), we identified 13 SNPs that reside within open chromatic regions (OCRs), each showing varying degrees of linkage disequilibrium (LD) with the sentinel SNP (**Figure 1**). Notably, rs3851179 itself does not resides within a OCR, but it is in high LD with rs10792832 (R^2^=0.99), which is located 765 bp away and colocalizes within a microglia-specific OCR and is reported to functionally disrupt a PU.1 transcription factor binding site. The PU.1 transcription factor is encoded by the gene *SPI1*, which is itself a strong AD risk gene (Kozlova et al., 2025).

**Figure 1.**
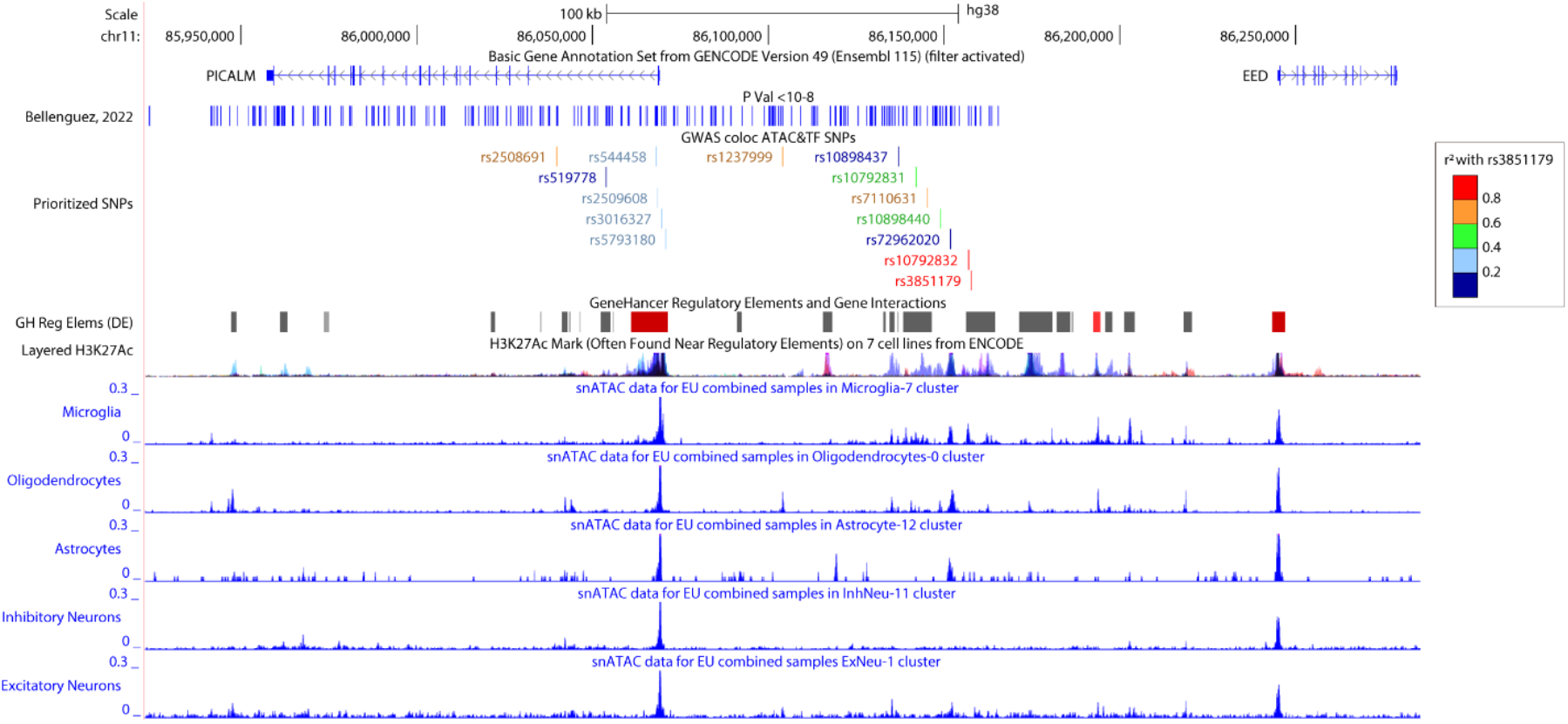
Prioritization of functional variants at *PICALM_EED* locus. The “Bellenguez, 2022” tracks display the 216 SNPs which are genome-wide significant in association with LOAD (Bellenguez et al., 2022). The “Prioritized SNPs” track displays the GWAS sentinel SNP and 13 candidate SNPs (color coded for LD degree with rs3851179), prioritized by co-localization on an OCR on the snATAC-seq peaks in each of the 5 major cell types: oligodendrocyte, microglia, astrocyte, inhibitory neuron and excitatory neuron are displayed in separate tracks (Celis et al., 2023).

### Cell-specific interactions between the GWAS sentinel SNP rs3851179 and surrounding chromosome region

To map potential causal target genes of the GWAS sentinel SNP rs3851179, we performed an eHiCA analysis using a 5 kb bait centered on rs3851179 (Bait #1), which also includes rs10792832. In the frontal cortex and microglia, the bait strongly interacts with the promoter and first intron of *PICALM* (**Figure 2**, orange highlight). In addition to this primary interaction, weaker interactions were observed between the bait and a cluster of CREs (CREe), defined by the presence of H3K27Ac marks in ENCODE (**Figure 2**, blue highlight). This CREe (chr11:86,192,334-86,203,859) is approximately 50kb away from the canonical TSS of *EED* and 130kb away from the TSS of *PICALM*. These interactions were not observed in iPSC-derived spheroids which are made up of oligodendrocytes, neurons, and astrocytes but which lack microglia (**Figure 2**) suggesting this is a microglia-specific interaction

**Figure 2.**
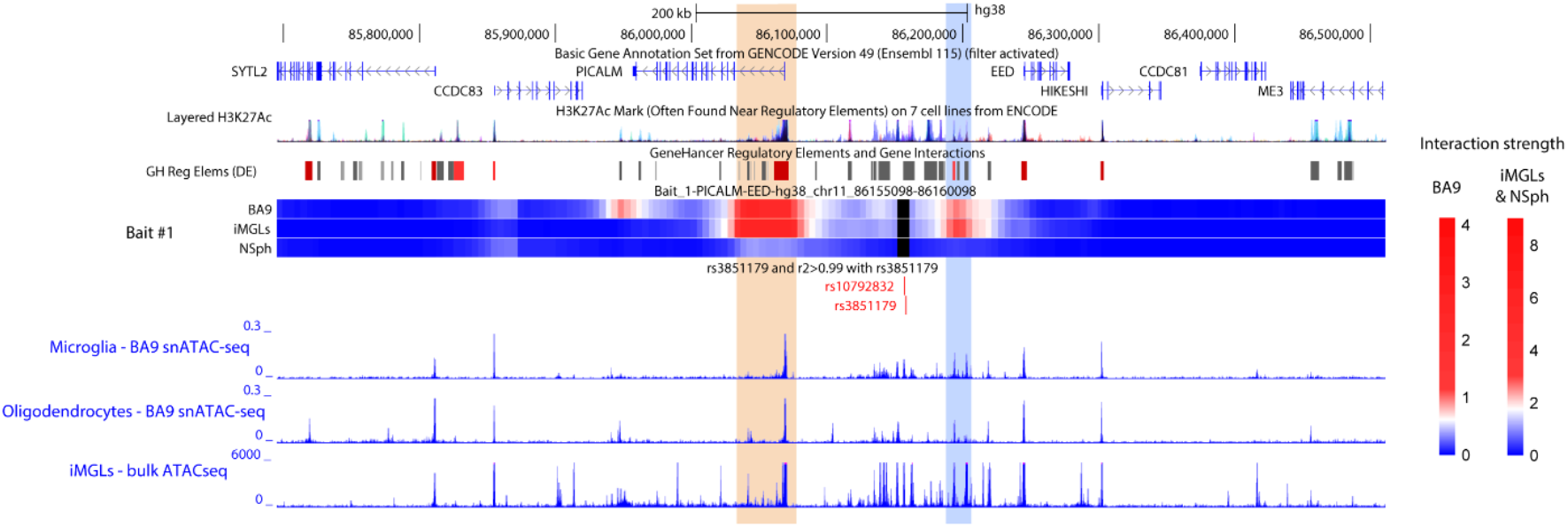
GWAS sentinel SNP bait predominantly interacts with the PICALM promoter and first intron. Heatmaps from eHi-C analysis show chromatin interactions using the bait (black bars) centered on rs3851179 and rs10792832. Color scales (right) indicate interaction strength in BA9 tissue and iPSC-derived microglia (iMGLs) or neural spheroids (NSph): increased red denotes stronger interaction, whereas blue indicates minimal or no interaction. Chromatin contacts between each bait and surrounding regions are displayed across an 0.8-Mb genomic window. No significant interactions were detected outside this interval. Chromatin accessibility data are derived from snATAC-seq profiles of microglia and oligodendrocytes (Celis et al., 2023), and bulk ATAC-seq from iMGLs (Moura et al., 2025).

### Chromatin interactions of other prioritized variants at the *PICALM/EED* locus

To comprehensively map potential causal variants and their target genes at the *PICALM/EED* locus, we expanded the application of eHiCA to other prioritized genetic variants in the region that are associated with LOAD (**Figure 1**). In addition to Bait #1, representing the sentinel SNP and rs10792832, eight additional baits were chosen (Bait #2-9) that lie upstream of Bait #1 and cover the remaining prioritized SNPs located in OCRs in this locus. Four prioritized SNPs reside in the proximal promoter of *PICALM*, spanning a 2.7 kb region (SNPs in light blue in the “GWAS coloc ATAC&TF SNPs” track in **Figure 1**) and are captured by one bait (Bait #4). This block of promoter SNPs is in high LD with each other (r^2^ > 0.9) but have limited LD (r^2^ = 0.32∼0.33) with the sentinel SNP rs3851179. Three prioritized SNPs (rs519778, rs10898437, and rs72962020) are in low LD with rs3851179 (r^2^ = 0.09∼0.13) and these were captured by three independent baits (Bait #3, #6, and #10, respectively). The remainder of the prioritized SNPs have moderate LD with the GWAS sentinel SNP rs3851179 (r^2^=0.44∼0.74) and were captured by five distinct baits (Bait #2, #5, #7, #8, and #9).

The bait interactions are shown in **Figure 3**. Baits #2-5 comprising prioritized SNPs in the 80 kb region in the first intronic region and promoter of the *PICALM* display a similar interaction profile, showing stronger interactions with the CREe cluster and the *EED* promoter region. Baits #6-9 overlap with a large enhancer cluster and share similar interaction patterns: strong interactions with the *PICALM* promoter and weaker interactions with the CREe cluster.

**Figure 3.**
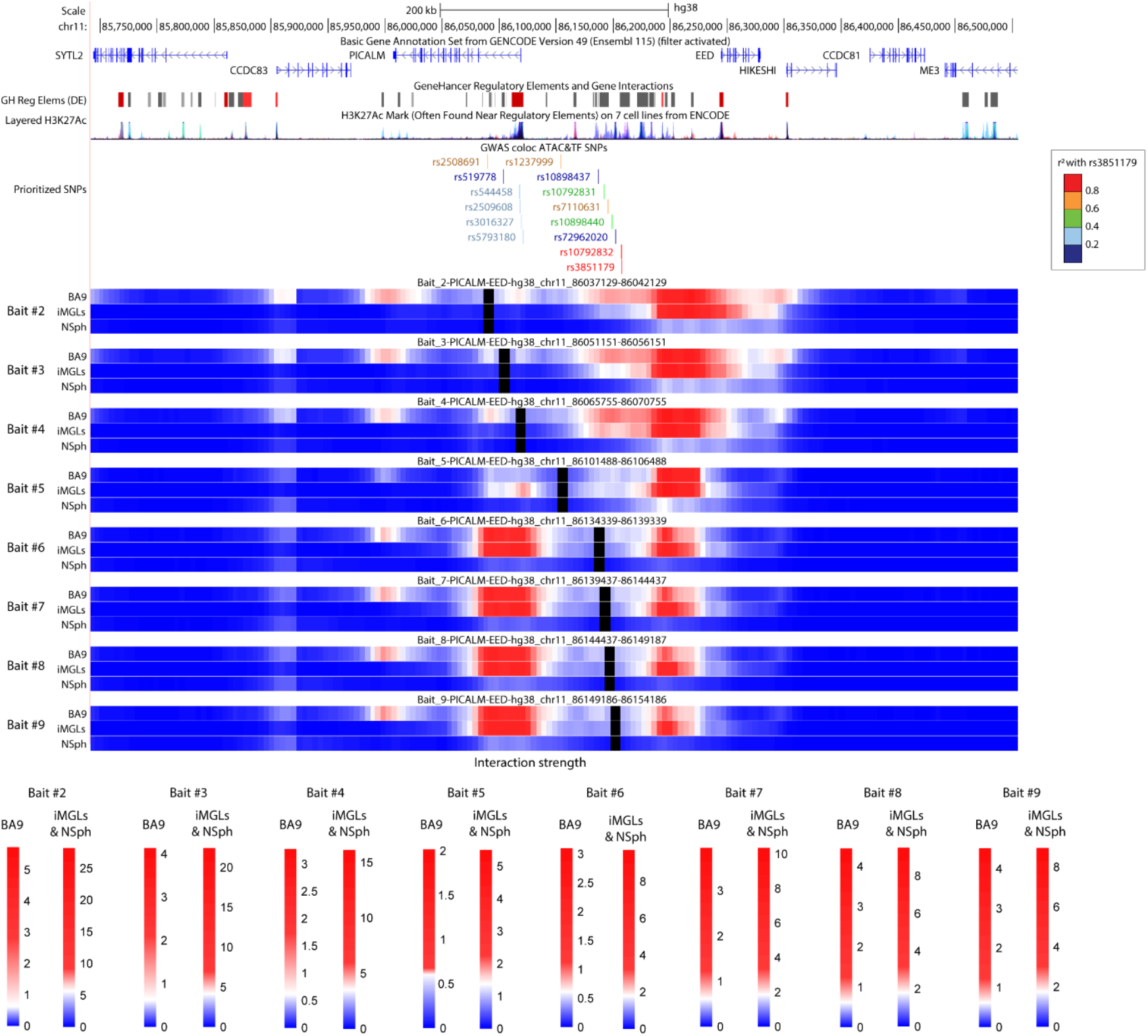
eHiCA of prioritized SNPs at *PICALM*_EED locus. Eight additional baits were designed to cover prioritized genetic variants that are associated with LOAD and located in OCRs. Baits (#2-9) were numbered based on the distance from rs3851179 sentinel SNP. Heatmaps from eHi-C analysis show chromatin interactions using the bait (black bars) including the respective SNPs. Color scales (bottom) indicate interaction strength in BA9 tissue and iPSC-derived microglia (iMGLs) or neural spheroids (NSph): increased red denotes stronger interaction, whereas blue indicates minimal or no interaction. Chromatin contacts between each bait and surrounding regions are displayed across an 0.8-Mb genomic window. No significant interactions were detected outside this interval.

### Reciprocal Chromatin Interactions at *PICALM* and *EED* Promoters

To understand the regulatory chromatin regions that interact with the promoters of *PICALM* and *EED* and the CREe locus, we performed eHiCA using the promoters of each gene as baits and four baits across the CREe (**Figure 4**). As stated before, *PICALM* promoter (Bait #4) displays reciprocal interactions with CREe as well as the promoter of *EED*. The *EED* promoter interacts with *PICALM* gene body and promoter region but not with the CREe. It also interacts with the promoter region of Coiled-Coil Domain Containing 83 gene (*CCDC83)* in brain and iMGLs. In the spheroids, the *EED* promoter has relatively strong interaction with the first intronic region of *PICALM* and weaker interaction with the *CCDC83* promoter. *CCDC83* is normally testis-specific but has been reported as a biomarker in several cancers (Kim et al., 2021; Song et al., 2012).

**Figure 4.**
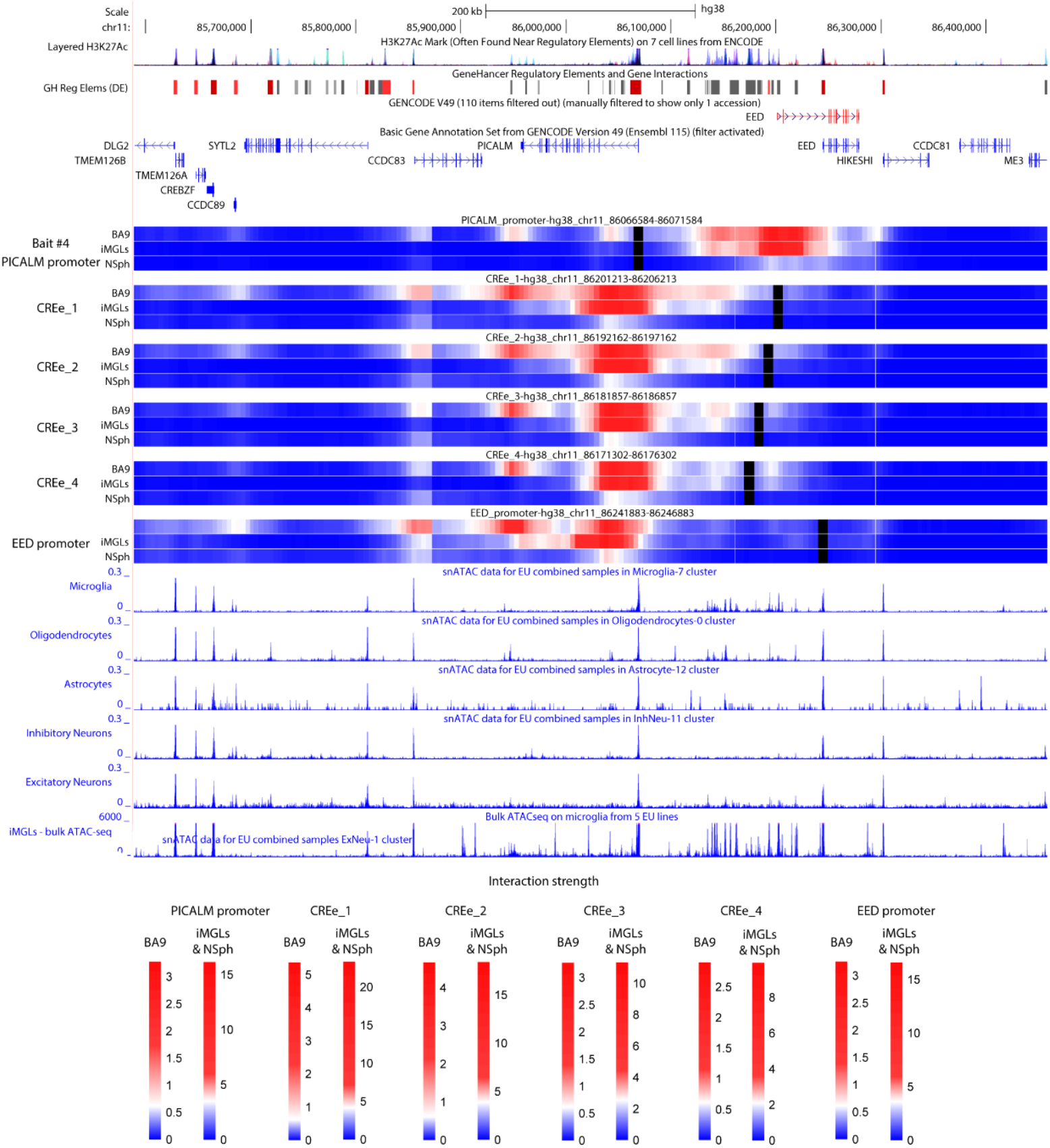
Reciprocal eHiCA of regulatory elements in *PICALM/EED* locus. Heatmaps of eHiCA analysis using the bait represented by the black bar at the promoter of *PICALM (or Bait #4)*, CREe, and promoter of *EED*. Color scales (bottom) indicate interaction strength in BA9 tissue and iPSC-derived microglia (iMGLs) or neural spheroids (NSph): increased red denotes stronger interaction, whereas blue indicates minimal or no interaction. Chromatin contacts between each bait and surrounding regions are displayed across an 0.9-Mb genomic window. No significant interactions were detected outside this interval.

### RNA sequencing

In both frontal cortex and iMGLs transcriptomic datasets, *PICALM* showed substantially higher expression than *EED*, with different patterns of cellular specificity. In snRNA-seq frontal cortex, *PICALM* expression was an order of magnitude higher than *EED* and was most enriched in microglia and endothelial cells, whereas *EED* expression was uniformly low and detected primarily in excitatory neurons, but also in microglia (Figure 5). The same pattern was detected in iMGLs (Figure 6 A).

**Figure 5.**
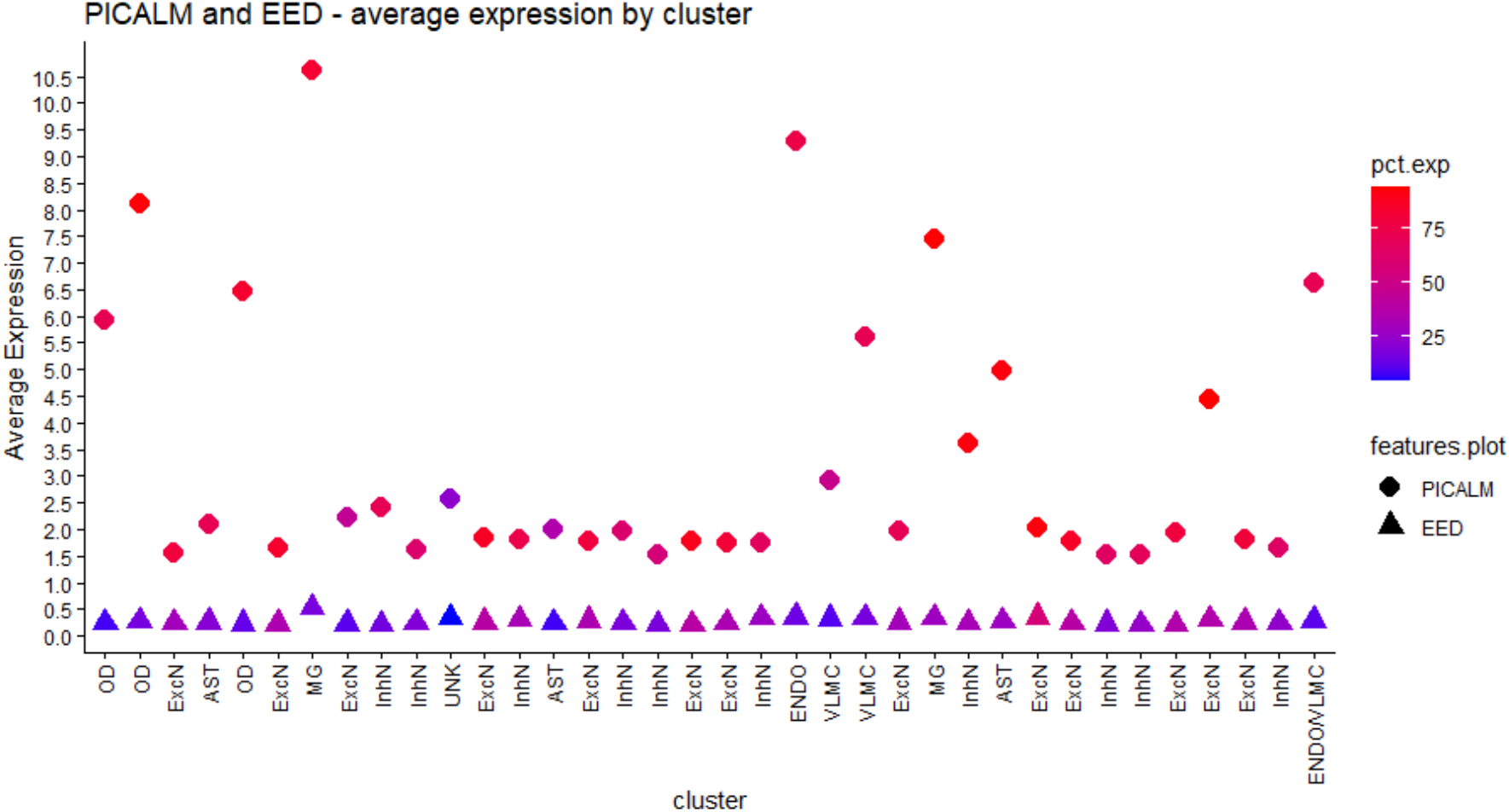
Average expression of *PICALM* and *EED*. Average expression levels of *PICALM* and *EED* in snRNA-seq in frontal cortex (BA9) (Griswold et al., 2021). OD: oligodendrocytes; ExcN: excitatory neurons; AST: astrocytes; MG: microglia; InhN: inhibitory neurons; ENDO: endothelial cells; VLMC: vascular leptomeningeal cells.

**Figure 6.**
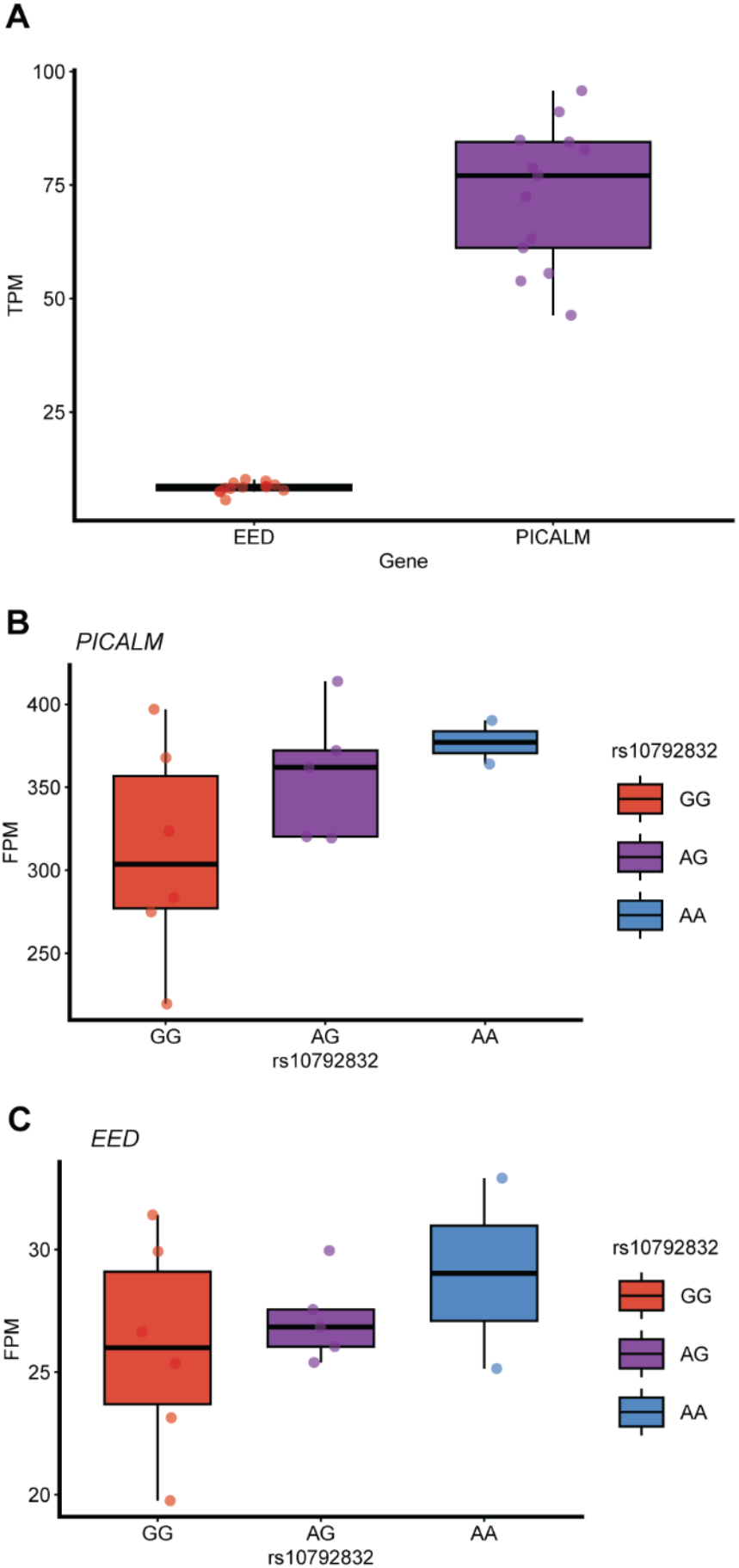
*PICALM* and *EED* expression in iMGLs. (A) *PICALM* and *EED* transcript abundance measured as Transcripts per Million (TPM) in induced microglia-like cells (iMGLs). TPM values adjusts for both gene length and sequencing depth, enabling direct comparison of transcript abundance across genes and samples. (B) *PICALM* expression quantified as Fragments per Million (FPM) across rs10792832 genotype groups (GG, AG, AA). FPM provides a measure of relative expression that accounts for sequencing depth but not transcript length. (C) *EED* expression quantified as FPM across rs10792832 genotype groups (GG, AG, AA), calculated as described in panel B. Boxplots summarize group-level distributions, and individual points represent sample-level expression.

To further test whether the presence of the risk haplotype at rs3851179 and rs10792832 alters gene expression, we examined the expression of *PICALM* in iMGLs stratified by genotype at rs3851179/rs10792832. Consistent with the microglia-specific regulatory mechanism suggested by PU.1 occupancy (Kozlova et al., 2025), carriers of the risk allele (G) showed reduced *PICALM* expression, whereas the rs10792832 A/A displayed the highest expression. This allele-dependent reduction in *PICALM* was not observed in *EED* (Figure 6 B and C).

Although our eHi-C analyses revealed a strong chromatin interaction between several baits (Baits #2–5) - including the PICALM promoter region - and a putative upstream promoter annotated as an alternative *EED* transcript (ENST00000707108.1), we found no transcriptional evidence for this isoform (Figure 4, GENCODE V49 filtered gene track, red). This transcript was absent from our bulk RNA-seq of iPSC-derived cell cultures (iMGLs and spheroids), absent from brain snRNA-seq, and was similarly undetectable in GTEx, GEO, and other publicly available datasets. In contrast, canonical *PICALM* and *EED* transcripts were readily detected in both brain and iMGL RNA-seq.

## Discussion

We demonstrate that the sentinel SNP (rs3851179) and the tightly linked variant rs10792832 interact directly with the *PICALM* promoter and a CRE cluster in microglia. Importantly, our data supports the finding of Kozlova et al. (2025) who demonstrated that the risk allele of rs10792832, which resides within a predicted PU.1 binding site, disrupts the PU.1 consensus motif and reduces PU.1 binding. Our RNA-seq data corroborates these findings supporting the association of the risk genotype (rs10792832 G/G) with decreased *PICALM* expression in microglia. Together, these data strongly supports that the association of rs3851179 with risk for AD is due to the direct interaction of rs10792832 with the PICALM promoter leading to reduced PICALM expression.

Understanding the target genes driving the pathobiology from genomic loci identified through genome-wide associations studies have been a challenge for many years and has hampered a full understanding of the genetic architecture underlying these conditions. Applying this strategy to the *PICALM/EED* region revealed a complex interaction matrix, where multiple baits designed to comprise SNPs with different levels of LD with the sentinel SNP reported by Bellenguez et al. (2022) interact with multiple loci. The eHiCA interactions anchored at the sentinel SNP rs3851179 (Bait #1, also comprising rs10792832) had a strong interaction with the *PICALM* promoter, strongly pointing to *PICALM* as the main effector gene. However, the sentinel SNP also has a weaker interaction with a cluster of CREe, but not with the designated *EED* promoter, indicating that additional genomic regions, particularly the CREe cluster, may be involved in the GWAS association. Notably, the analysis also showed that the regulatory landscape of this region is cell-type specific, since the interactions between *PICALM* and the distal CREe cluster were observed in iPSC-derived microglia (iMGLs), whereas these interactions were largely absent in iPSC-derived spheroids lacking microglia, supporting a microglia-specific regulatory mechanism.

Promoter-anchored analyses show that the *PICALM* promoter interacts not only with the CREe cluster but also with the *EED* promoter, suggesting a shared chromatin domain that may be dynamically configured across cell types. In contrast, *EED* promoter interactions are more restricted, engaging with *PICALM* but not with the CREe. The interaction of the promoter of *EED* with *PICALM* could reflect chromatin proximity driven by overall locus structure rather than direct co-regulation, especially given that *EED* expression is largely neuronal (in our dataset and the Allen transcriptome explorer dataset) and not enriched in microglia, where *PICALM* is most highly expressed. This supports a model in which *PICALM* is the primary effector gene of the AD-associated locus in microglia, while *EED* and nearby genes such as *CCDC83* may participate in alternative or secondary regulatory contexts.

Understanding the complex regulatory landscape between GWAS identified loci and the target genes responsible for contributing to the disease process remains a priority in the study of complex human disorders. This has been complicated by two elements; the haplotype structure of the genome and the increasing cohort size used for GWAS, which increases the power to identify less prominent associations. This appears to be the case with the *PICALM/EED* locus, where over 200 SNPs reach genome-wide association, bringing loci at lower R^2^ in the haplotype to the forefront. Therefore, it becomes important in elucidating the difference between variants present in the haplotype due to inherited co-segregation within a haplotype with those variants driving the association due to their biological function. A haplotype represents DNA variants traveling together across generations, but such linkage disequilibrium does not necessarily imply that the variants within the haplotype share regulatory functions or participate in similar chromatin interactions. The average genomic haplotype size is estimated to be about 9∼15kb in the African genome and 18∼25 kb in the European genome (Chimusa et al., 2015). Since our eHICA approach relies on 5 kb baits which are much smaller than the average haplotype sizes, it allows us to partially deconvolve these haplotype blocks, assigning different segments to distinct chromatin interaction bins, and potentially perform different functions. Also, we can see that different interactions are seen in different cell types. This highlights the need to focus on cell-type-specific OCRs, such as those identified in microglia.

In summary, the chromatin interactions demonstrate that *PICALM* is the primary gene tied to the GWAS association for AD at this locus and strongly supports that the association of rs3851179 with risk for AD is due to the direct interaction of rs10792832 with the PICALM promoter of PICALM leading to reduced PICALM expression. The CREe cluster 5’ to *EED* is also highly interactive and may modulate *PICALM* promoter activity through long-range looping, particularly in microglia. Finally, we believe the eHiCA approach will provide important insight into GWAS associations and help define the target gene(s) with in these association regions providing important information on disease mechanisms and highlighting putative, biologically relevant therapeutic targets.

## Supporting information

Samples Used

## Acknowledgements

We acknowledge the Center for Genome Technology (CGT) from the John P. Hussman Institute for Human Genomics (HIHG) from the University of Miami, Miller School of Medicine for the genomic and data analyses. We thank and express our gratitude to the numerous participants, researchers, and staff involved for their invaluable contributions to the present study. This study was supported by the National Institute on Aging (grant numbers U01-AG072579, RF1-AG059018, U01-AG066767, U01-AG052410, R56-AG072547, R01-AG070864, U19AG074865) the Hussman Foundation and the Alzheimer Association.

## Author Contributions

LW, FJ, DD, JIY and JMV contributed to the conception and design of the study. LBN, LW, W Xu, AR, SM, LL, XL, FR, KC, MG, DAB, SW, CG, TS, WKS, MAP, JIY, FL and KN contribute the acquisition and analysis of data, LBN, LW, DMD, JIY, AJG, FJ, KN, WX, and JMV contributed to drafting of manuscript or figures.

